# Measuring Genomic Data with Prefix-Free Parsing

**DOI:** 10.1101/2025.02.21.639270

**Authors:** Simone Lucà, Francesco Masillo, Zsuzsanna Lipták

**Author notes:** Corresponding author (Simone Lucà). Email addresses* (Francesco Masillo), (Zsuzsanna Lipták).

## Abstract

**Summary:** Prefix-free parsing [Boucher et al., Alg. Mol. Biol., 2019] is a highly effective heuristic for computing text indexes for very large amounts of biological data. The algorithm constructs a data structure, the prefix-free parse (PFP) of the input, consisting of a dictionary and a parse, which is then used to speed up computation of the final index. In this paper, we study the *size* of the PFP, which we refer to as *π*, and show that it is a powerful tool in its own right. To show this, we present two use cases. We first study the application of *π* as a *repetitiveness measure* of the input text, and compare it to other currently used repetitiveness measures, including *z* (the number of Lempel-Ziv phrases), *r* (the number of runs of the Burrows-Wheeler Transform), and *δ* (the text’s substring complexity). We then turn to the use of *π* as a measure for *pangenome openness*. In both applications, our results are similar to existing measures, but our tool, in almost all cases, is more efficient than those computing the other measures, both in terms of time and space, sometimes by an order of magnitude. We close the paper with a detailed systematic study of the parameter choice for PFP (window size *w* and modulus *p*). This gives rise to interesting open questions.

**Availability and implementation:** The source code is available at https://github.com/simolucaa/piPFP. The accession codes for all the datasets used and the raw results are available at https://github.com/simolucaa/piPFP_experiments.

## 1. Introduction

With the availability of unprecedented amounts of genomic data today, it is becoming increasingly important to be able to store, handle, and process this data efficiently (Pan-Genomics Consortium, 2018). One central issue is that datasets, in addition to their size, can have different properties, such as number of sequences, length of individual sequences, or level of repetitiveness. Current software is frequently tailor-made for specific types of data, taking advantage of certain properties of the input. Therefore, it is paramount that users be able to test their data’s properties.

One such central property is *repetitiveness*. Intuitively, this is a measure of how much of the sequence data occurs more than once in the input. This concept is intimately related to our ability to store the data using compressed text indexes, which require little storage space while at the same time enabling efficient string processing tasks such as pattern matching (Navarro, 2022b). To date there is no generally accepted measure of repetitiveness (Navarro, 2022a), and different measures are being used, such as the number *z* of the phrases of the Lempel-Ziv 77 parse (Lempel and Ziv, 1976; Ziv and Lempel, 1977), the number *r* of runs of the Burrows-Wheeler Transform (BWT) (Burrows and Wheeler, 1994), or the more recent *δ*-measure (Raskhodnikova et al., 2013) (also referred to as substring complexity).

Another property of interest is *pangenome openness* (Tettelin et al., 2008). A pangenome is a set of genomes of individual of the same species. Pangenome openness essentially measures the amount of additional data necessary to have complete information about a species, given a dataset of individual genomes (i.e., a pangenome). Several methods for measuring pangenome openness were introduced recently (Parmigiani et al., 2024; Bonnie et al., 2024; Page et al., 2015; Jonkheer et al., 2022; Chaudhari et al., 2016).

In this paper, we introduce a new method of measuring genomic data, based on prefix-free parsing, and show that it can be successfully applied in both of these areas: as a repetitiveness measure and for measuring pangenome openness.

*Prefix-free parsing* (Boucher et al., 2019) (PFP) is a highly effective heuristic for computing text indexes for very large amounts of biological data. The algorithm computes a data structure, the prefix-free parse (PFP) of the input, consisting of a dictionary and a parse, which is then used to speed up computation of the final index. Originally introduced as a preprocessing step to speed up the computation of the Burrows-Wheeler Transform (BWT) of very large genomic sequence data, it has since been extended to computing other data structures, as well. These include the suffix array (Kuhnle et al., 2020); the extended BWT (Boucher et al., 2021a, 2024) of Mantaci et al. (Mantaci et al., 2007); and the XBWT (Gagie et al., 2021), an extension of the BWT to labelled trees. PFP has also been introduced as a text index in its own right (Boucher et al., 2021b; Oliva et al., 2022), or as a way to enhance the FM-index to boost query performance (Hong et al., 2024); and a recursive variant for BWT construction was proposed (Oliva et al., 2023) in order to further scale the computation with huge amounts of data. PFP has also shown competitive results for building grammars (Gagie et al., 2019; Kim et al., 2024), computing the Lempel-Ziv factorization (Hong et al., 2023), and computing the smallest suffixient set (Cenzato et al., 2025). Other applications of PFP for constructing pangenome-sized data structures include building the *r*-index of Gagie et al. (Gagie et al., 2020) augmented with thresholds (Bannai et al., 2020). This combination of data structures enable the computation of matching statistics, which in turn allows finding Maximal Exact Matches (MEMs). MEMs are a crucial ingredient for speeding up the alignment phase of sets of reads to a reference genome. Recent tools, such as `MONI` (Rossi et al., 2022), `PHONI` (Boucher et al., 2021c), `AUG-PHONI` (Martínez-Guardiola et al., 2023), `LAZY-PHONI` (Goga et al., 2024), `SPUMONI` (Ahmed et al., 2021), `SPUMONI2` (Ahmed et al., 2023), `SIGMONI` (Shivakumar et al., 2024), `MOVI` (Za-keri et al., 2024), all use PFP as preprocessing step for building the underlying *r*-index. Latest work includes the use of PFP for computing multiple sequence alignments (Olbrich et al., 2025) and for merging very large BWTs (Díaz-Domínguez et al., 2025).

Here, we explore the power of PFP for measuring certain properties of the input data. We introduce the measure *π*, the *size* of the PFP data structure, and show that it is a powerful tool in its own right. Our experiments in fact indicate that *π* might be a good candidate for a new measure both as a repetitiveness measure, and for measuring pangenome openness. It gives similar results regarding repetitiveness as the established repetitiveness measures *z, r*, and *δ*. When used to decide pangenome openness, on almost all of the benchmark datasets, classification using *π* gives the same result as recent competitor measures.

We also present a tool, `piPFP`, which computes *π*. Our experiments show that `piPFP` is highly efficient both as regards computation time and memory requirements, and outperforms other methods sometimes by orders of magnitude. We believe that using *π* as an alternative method for measuring these properties can be beneficial to users, as it requires little computational resources and has similar accuracy as other, more complex and/or slower, methods.

Finally, we turn our attention to parameter choice in PFP. The PFP algorithm has two user-defined parameters, the window size *w* and the modulus *p*. In the original paper (Boucher et al., 2019), a brief study of parameter choice was presented, on the basis of which the default parameters of the accompanying tool were chosen. Another parameter comparison was included in the PhD thesis (Oliva, 2023) (see Sections 3.1 and 3.3 for detailed discussions). Here, we explore these parameters in depth, concluding that for certain applications, very small window sizes may not be appropriate. Our study on the parameter choices leads to several open problems of a theoretical nature. The rest of the paper is organized as follows. In Section 2, we give the necessary technical background and present our experimental setup. Section 3 contains the experimental results: as a repetitiveness measure, for pangenome openness, and exploring parameter choice for PFP. We close with a discussion and open problems in Section 4.

## 2. Materials and Methods

In this section, we first give the necessary background on prefix-free parsing (Sec. 2.1), the repetitiveness measures *z, r*, and *δ* (Sec. 2.2), and pangenome openness (Sec. 2.4). This is followed by the introduction of our PFP-based measure (Sec. 2.5), the presentation of the datasets used in our experiments (Sec. 2.6), and details of the implementation (Sec. 2.7).

### 2.1. Prefix-Free Parsing

Prefix-free parsing (*PFP*), introduced in (Boucher et al., 2019), is a technique that processes a string *T* of length *n* using two integers, *w* and *p*, both greater than 1. The method produces a parse of *T* consisting of overlapping phrases, each uniquely stored in a dictionary denoted as *D*. The resulting parse *P* is represented as a meta-string, where each phrase is replaced by its rank in the lexicographically sorted dictionary. We refer to the prefix-free parse of *T* as *PFP*(*T*), consisting of the dictionary *D* and the parse *P*. Next, we give an outline of this method.

The PFP process begins by appending *w* occurrences of a special symbol #, which is smaller than all characters and does not appear elsewhere in the string, to both the beginning and end of *T*. We refer to the resulting string as *T* ^′^ = #^*w*^*T* #^*w*^. Next, we define a set of *trigger strings, E*, that will guide the parsing of *T*, as follows: We compute the Karp-Rabin hash *H*_*q*_(*t*) for all substrings of length *w* in *T* ^′^ using a large prime *q*. A substring *t* = *T* ^′^[*i*..*i* + *w* − 1] is then in *E* if *H*_*p*_(*t*) = *H*_*q*_(*t*) mod *p* = 0, or if *t* = #^*w*^. This can be computed efficiently via the well-known rolling hash algorithm of Karp and Rabin (Karp and Rabin, 1987). These trigger strings are then used to parse *T* ^′^, as follows.

The dictionary *D* for PFP consists of substrings of *T* ^′^ that start with a trigger string and end with the next trigger string in *T* ^′^. These substrings, referred to as *phrases*, can be identified by scanning *T* ^′^ and adding a new phrase to *D* whenever the next trigger string is encountered. Once the dictionary *D* is built, it is sorted lexicographically. Using the sorted dictionary *D* and the input string *T*, we generate the parse *P*, which consists of these overlapping phrases from *D*. Each phrase overlaps with the next by *w* characters, corresponding to a trigger string. This parsing process creates an array of integers in *P*, where each integer is the rank of the corresponding phrase in *D*. With *D* and *P*, the original string *T* can be reconstructed.

A set of strings is called *prefix-free* if it does not contain any two strings which are one a proper prefix of the other. It can be shown that the set of the suffixes of phrases (i.e., elements of the dictionary *D*) are prefix-free, as formalized in the following lemma:

#### Lemma 1

((Boucher et al., 2019), **Lemma 1)**. *For a string T and its prefix-free parse PFP*(*T*), *which consists of a dictionary D and a parse P*, *the set S of suffixes of phrases in D of length at least w forms a prefix-free set*.

This is a crucial property of PFP, necessary for the fast building of text indexes such as the BWT. It was shown in (Boucher et al., 2019) how to construct the Burrows-Wheeler Transform (*BWT*) (Burrows and Wheeler, 1994) using space and time proportional to the size of the dictionary *D* and parse *P*. Later, in (Kuhnle et al., 2020), it was shown how to compute the suffix array samples at run-boundaries while computing the *BWT* to produce the *r*-index (Gagie et al., 2020) within the same time and space bounds.

#### Example 1.

*Let the parameters be w* = 2 *and p* = 11. *(We ignore q here, as it is a large prime, and has therefore no impact on the computation, i.e*., *H*_*q*_(*u*) = *u for short strings u.) Using b* = 256 *and the ASCII-256 encoding, where* `A` = 65, `C` = 67, `G` = 71, *and* `T` = 84, *the only two w-length words that map to* 0 *are* `GT` *and* `TC`:

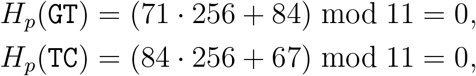

*and therefore, E* = {##, `GT, TC`}. *Consider the string*

*T* = TCCGGGCTGGCATCTGCCGTAAAGGTGCGGATATCCGGGCTGGCATCA

*T* ^′^ = ##TCCGGGCTGGCATCTGCCGTAAAGGTGCGGATATCCGGGCTGGCATCA##

*With E* = {##, `GT`, `TC`}, *we compute the corresponding lexicographically sorted dictionary: D* = {[0: ##`TC`], [1: GTAAAGGT], 2: GTGCGGATATC], [3: `TCA`##], [4: TCCGGGCTGGCATC], [5: TCTGCCGT]}.

*The parse that is obtained from T* ^′^ *and D is P* = [0, 4, 5, 1, 2, 4, 3].

### 2.2. Repetitiveness measures

The study of string repetitiveness focuses on quantifying the complexity and redundancy present in a string. For relationships among the wide spectrum of repetitiveness measures, we point the reader to the comprehensive survey (Navarro, 2022a).

#### 2.2.1. Number of phrases in the Lempel-Ziv factorization

The Lempel-Ziv compression algorithms (Lempel and Ziv, 1976; Ziv and Lempel, 1977, 1978) are fundamental to understanding repetitiveness in strings. In particular, the LZ77 compression algorithm (Lempel and Ziv, 1976; Ziv and Lempel, 1977) parses a string *T* into a sequence of phrases, where each phrase is either a new character or a substring of maximum length that appears also earlier in the text. The number of phrases, denoted as *z*, is a good measure of repetitiveness, since highly repetitive strings tend to have fewer phrases.

Lempel-Ziv parsing is significant not only as a compression technique but also as a way to define theoretical bounds for other measures of repetitiveness. It captures the idea of redundancy in a string by representing repeated patterns succinctly.

#### 2.2.2. Number of runs of the Burrows-Wheeler Transform

The Burrows-Wheeler Transform (*BWT*) (Burrows and Wheeler, 1994) is another method which is widely used in string processing and compression. Given a string *T*, the *BWT* is computed by sorting lexicographically the set of suffixes of *T* and then concatenating the preceding character of the suffixes. A well-known property of the *BWT* is that repetitive strings tend to produce long runs of identical characters in the transformed string. The number of runs, denoted as *r*, can thus be used as a measure of repetitiveness, since smaller values of *r* indicate higher levels of repetitiveness in the original string.

The number of runs *r* is closely related to the compressibility of *T* and can be exploited for building text indexes, such as the FM-index (Ferragina and Manzini, 2000), or the more recent *r*-index (Gagie et al., 2020) or extended *r*-index (Boucher et al., 2024).

### 2.3. Substring complexity

The measure *δ*, known as *substring complexity* (Raskhodnikova et al., 2013), quantifies the repetitiveness of a string based on the normalized number of distinct substrings of fixed length *k*, maximized over all possible *k*. Formally, for a string *T* of length *n*, the measure *δ* is defined as:

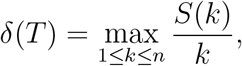

where *S*(*k*) is the number of distinct *k*-length substrings of *T*.

The measure *δ* is known to lower-bound both *z* and *r* for every text *T* (Kociumaka et al., 2023).

### 2.4. Pangenome openness

The term pangenome openness (Tettelin et al., 2008) refers to a method to estimate a pangenome’s total size, given a set of genomes of individuals from the same species. This estimation requires calculating the pangenome growth, which monitors the number of distinct attributes observed as a function of the number of entities considered. Heap’s Law (Heaps, 1978), known in the information retrieval field, can be used to describe this phenomenon as follows: the rate at which new attributes are found decreases as one considers more and more entities, as this rate is proportional to *kN*^*γ*−1^ = *kN* ^−*α*^, where *α* = 1 − *γ* and *k* are constants that depend on the particular text, and *N* is the total number of attributes. In other words, as sampling proceeds, discovering a new attribute becomes increasingly harder (Tettelin et al., 2008; Heaps, 1978; Parmigiani et al., 2024). In pangenomics, the entities are the genomes, and the attributes can be genes, open reading frames (ORFs), or genome intervals of fixed size (e.g., *k*-mers (Parmigiani et al., 2024)). The parameter *α* is used to classify a genome as either *closed* or *open*:

- If *α <* 1, the pangenome is classified as *open*. This means that the number of attributes discovered increases significantly with the addition of new genomes. This indicates a high level of genomic diversity, suggesting that each newly sequenced genome contributes unique genes.
- If *α >* 1, the pangenome is classified as *closed*. This indicates that the number of new attributes reaches a plateau as more genomes are added to the set.

This definition of pangenome growth, however, makes *α* dependent on the order in which the genomes are added. To address this issue, one can average over all possible permutations; however, this is prohibitive, even for a small number of genomes. One solution is to estimate the growth by selecting a subset of all possible permutations (Tettelin et al., 2008; Bonnie et al., 2024). We will instead employ a solution presented by Parmigiani et al. (Parmigiani et al., 2024), which is based on a commonly used method in ecology (Heck Jr et al., 1975). In the following, we follow the definitions given in (Parmigiani et al., 2024). (Note that their definition of openness differs slightly from the original one given in Tettelin et al. (Tettelin et al., 2008), as explained in (Parmigiani et al., 2024, p. 5).)

Given *n* entities *G*_1_, …, *G*_*n*_, we define the average total cardinality *f*_*tot*_ (*m*), for 1 ≤ *m* ≤ *n*, of the union of *m* of these entities, and the average number *f*_*new*_(*m*) of new attributes that are added when adding an *m*th entity to *m* − 1 entities, as follows:

#### Definition 1.

*Let* U *be a universe (of* attributes*) and* G = {*G*_1_, …, *G*_*n*_} *a set of* entities, *i.e. G*_*i*_ ⊆ U *for* 1 ≤ *i* ≤ *m. Then*

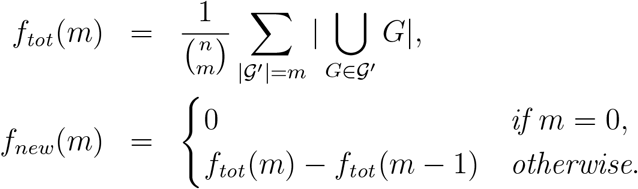

*We also write f*_*tot*_ *for f*_*tot*_ (*n*) *and f*_*new*_ *for f*_*new*_(*n*).

It is known (Parmigiani et al., 2024) that *f*_*new*_ follows Heap’s Law, therefore it is possible to fit *kN* ^−*α*^ on *f*_*new*_ and derive the parameter *α*.

### 2.5. How we use PFP for measuring genomic data

To evaluate the repetitiveness of a sequence, we rely on the fact that PFP phrases in the dictionary are unique. Ideally, a more repetitive sequence will result in fewer phrases in the dictionary, with the parameters *w* and *p* influencing the number of phrases. In order to better study this question, we introduce a measure of the size of the PFP which encompasses the two parameters *w* and *p*, as follows:

#### Definition 2.

*Let w, p be two integers, where w* ≥ 4 *and p* ≥ 10. *Given a text T*, *let D*_*w*,*p*_ *and P*_*w*,*p*_ *be the dictionary and parse, respectively, of the PFP using window size w and modulus p. We define*

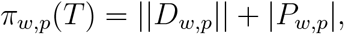

*where* || *D*_*w*,*p*_|| *denotes the total length of the phrases in D*_*w*,*p*_ *and* |*P*_*w*,*p*_| *the length of the parse. When T is clear from the context, we also write π*_*w*,*p*_ *for π*_*w*,*p*_(*T*).

#### Example 2.

*Consider the string*

*T* = TCCGGGCTGGCATCTGCCGTAAAGGTGCGGATATCCGGGCTGGCATCA

*and the parameters w* = 2 *and p* = 11.

{

*As shown in Example 1, the dictionary and parse of T with these parameters are as follows: D_2,11_ = [0: ##`TC`], [1: GTAAAGGT], [2: GTGCGGATATC], [3: `TCA`##], [4: TCCGGGCTGGCATC], [5: TCTGCCGT] and P*_2,11_ = [0, 4, 5, 1, 2, 4, 3]. *The total length of the phrases in D*_2,11_ *is* ||*D*_2,11_|| = 4+8+11+5+14+8 = 50, *and the length of the parse is* |*P*_2,11_| = 7. *Therefore, we have π*_2,11_(*T*) = ||*D*_2,11_|| + |*P*_2,11_| = 50 + 7 = 57.

To assess pangenome openness, we applied the approach of Parmigiani et al. (Parmigiani et al., 2024) described earlier, using PFP phrases as attributes. We compute a histogram *h*(*i*), which, for every *i*, counts how many PFP phrases are present in exactly *i* genomes. This histogram is then used to estimate *f*_*tot*_. Finally, we compute *f*_*new*_ from *f*_*tot*_ and fit the function *kN* ^−*α*^ to *f*_*new*_ to estimate *α*.

### 2.6. Datasets used

In this section, we present an overview of the datasets used in our experiments. We conducted three main sets of experiments, each using different datasets. These datasets are accessible through NCBI using the provided accession codes: https://github.com/simolucaa/piPFP_experiments. A summary of the datasets is presented in Table 1.

**Table 1:** Table summarizing the datasets used in the three main sets of experiments. Genome size is given in mega-bases (Mb), and *σ* is the size of the alphabet.

| | species | number of<br>genomes | average genome<br>size (Mb) | total size<br>(Mb) | $\sigma$ |
| --- | --- | --- | --- | --- | --- |
| 1 | *A. thaliana* | 21 | 136.64 | 2869.43 | 16 |
| 2 | *E. coli* | 120 | 5.74 | 665.63 | 8 |
| 3 | SARS-CoV-2 | 500 | 0.03 | 14.90 | 11 |
| 4 | *S. cerevisiae* | 28 | 12.04 | 337.23 | 9 |
| 5 | *S. enterica* | 60 | 4.80 | 288.25 | 14 |
| 6 | *H. sapiens* | 9 | 3081.83 | 26856.70 | 16 |
| 7 | *E. coli* | 7725 | 5.15 | 38414.10 | 15 |
| 8 | *B. aphidicola* | 35 | 0.61 | 20.45 | 5 |
| 9 | *B. cereus* | 83 | 5.65 | 452.63 | 13 |
| 10 | *C. botulinum* | 50 | 3.95 | 190.76 | 5 |
| 11 | *C. burnetii* | 13 | 2.06 | 25.86 | 4 |
| 12 | *C. jejuni* | 234 | 1.68 | 378.74 | 15 |
| 13 | *F. tularensis* | 57 | 1.90 | 104.65 | 5 |
| 14 | *H. pylori* | 234 | 1.63 | 368.63 | 15 |
| 15 | *P. marinus* | 9 | 1.73 | 15.04 | 4 |
| 16 | *R. palustris* | 8 | 5.38 | 41.53 | 4 |
| 17 | *S. pneumoniae* | 88 | 2.11 | 179.43 | 8 |
| 18 | *S. pyogenes* | 247 | 1.85 | 440.21 | 11 |
| 19 | *Y. pestis* | 56 | 4.71 | 254.80 | 6 |

#### Datasets used for π as a repetitiveness measure

To evaluate the effectiveness of *π* as a repetitiveness measure, we performed three experiments:

1. Comparison of *π* to other repetitiveness measures. For this comparison, we used the same 60 *Salmonella enterica* genomes as Bonnie et al. (Bonnie et al., 2024) (entry 5 in Table 1).
2. Comparison of *π* of real and random sequences. We generated random sequences (detail in Section 3.1) and compared their *π* values to those from 60 *S. enterica* genomes and 120 *Escherichia coli* genomes (see entries 5 and 2 in Table 1).
3. Performance assessment. For this experiment, we tested the tools on the same datasets as (2) and on larger datasets, namely 9 *Homo sapiens* genomes and 7725 *E. coli* genomes (entries 6 and 7 of Table 1).

#### Datasets used for pangenome openness

To evaluate our tool, we compared our results to those of other tools in two ways:

1. Prediction accuracy. We compared our predictions to those of other tools. For these experiments, we used the same dataset as Parmigiani et al. (Parmigiani et al., 2024), which comprises 12 prokaryotic genomes (see entries 8-19 of Table 1).
2. Performance assessment. For this experiment, we compared our tool to others. We used the same datasets as (1) and larger datasets, namely 9 *H. sapiens* genomes and 7725 *E. coli* (entries 6 and 7 of Table 1).

#### Datasets used for parameter choice for π

For this experiment, we selected five datasets to represent five different biological categories: a complex animal (*H. sapiens*), a plant (*Arabidopsis thaliana*), a simple eukaryote (*Saccharomyces cerevisiae*), two prokaryotic organisms (*S. enterica* and *E. coli*), and a virus (SARS-CoV-2). More information about these datasets can be found in Table 1 (entries 1-6).

### 2.7. Implementation details

Our tool `piPFP` is available at https://github.com/simolucaa/piPFP. We implemented both the computation of the repetitiveness measure *π*_*w*,*p*_ and the pangenome openness measure, building on the original implementation of PFP^1^.

We modified the original implementation in order to speed up the computation and to save working space. In the dictionary, we only store the hash of the phrases rather than the full explicit phrases. This is because we only need the number of distinct phrases and their length, not the actual phrases themselves. In practice, we maintain a hashmap that stores the hashes of the phrases as keys and their lengths as items. We also avoid storing the parse explicitly on disk by counting the total number of phrases found during computation, reducing the parse to a single integer counter. This is done in order to avoid costly I/O operations.

When using multiple threads, we also slightly deviate from the original implementation by using a concurrent hashmap, `growt`, from (Maier et al., 2019) instead of the standard implementation found in the C++ library. This allows faster parallel insertions of new entries in the hashmap without an explicit lock mechanism.

For the computation of pangenome openness (see Sections 2.4 and 2.5), we based our implementation on (Parmigiani et al., 2024), replacing the *k*-mers with PFP phrases. To enable fast computation across different threads, we assign to each thread a range of sequences to process and populate a thread-specific hashset. This thread-specific hashset stores only the hashes of the phrases. After having populated the hashset for one genome, there is a merging phase in which we merge the populated thread-specific hashset into a collective hashmap. This collective hashmap stores hashes of phrases as keys and the corresponding counter as items. The collective hashmap is then processed to output the final histogram after every genome is processed. In our program, we use the standard C++ library implementation for the hashsets, while we again use the `growt` implementation for the collective hashmap.

## 3. Experimental Results and Discussion

For all experiments presented in this section, we used a machine equipped with an Intel(R) Core(R) i9-11900 @ 2.50GHz (with turbo speed @ 5GHz) with 16 MB of cache and 8 cores (16 threads via hyperthreading) and 64 GB of DDR4-3200MHz RAM. The operating system was Ubuntu 22.04 LTS, and the compiler used was `g++` version 11.4.0. The code was compiled with options `-std=c++20 -O3 -march=native -ldl -pthread` enabled. Measurements regarding running times and space consumption were recorded using the built-in command of the operating system `/usr/bin/time -v`.

### 3.1. PFP as a repetitiveness measure

In this section, we evaluate our proposed repetitiveness measure, *π*_*w*,*p*_, through three sets of experiments. First, we compare *π*_*w*,*p*_ with the established repetitiveness measures *r, z*, and *δ*. Previous work has already compared the working space of PFP with *r*. In particular, Oliva (Oliva, 2023) found a linear correlation between the size of the PFP and *r* in one set of highly repetitive data. Additionally, Lucà (Lucà, 2023) studied how the degree of sequence repetitiveness influenced the sizes of *D* and *P* across different parameter combinations in various real and artificial datasets, and compared ||*D*|| to *r*. Second, we examine the behavior of *π*_*w*,*p*_ on real and randomly generated sequences. Third, we evaluate the performance of our tool and other tools for different measures, using both small and large datasets.

In the first experiment, we compare *π*_*w*,*p*_ (computed with `piPFP`) to the repetitiveness measures *z* (computed with `PFP_LZ77`), *r* (`optimalBWT`), and *δ* (`substring-complexity`, exact version). We ran `piPFP` with the default parameters *w* = 10 and *p* = 100, since *π* does not vary significantly with *w >* 10 and *p >* 50 (see Fig. 7). We used the dataset provided in (Bonnie et al., 2024), as detailed in Sec. 2.6 and Table 1.

Following the procedure of (Bonnie et al., 2024), we chose an arbitrary order of the input set of genomes. Then we computed the repetitiveness measures *z, r, δ*, and *π*_10,100_ on increasingly larger subsets of genomes, from 1 to 60, and applied 0-1 normalization for each measure individually:

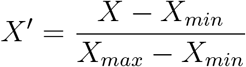

where *X* is the original value, *X*_*min*_ and *X*_*max*_ are the minimum and maximum values of the measure, and *X*^′^ is the normalized value.

We report the results in Figure 1. As we can observe, *π*_10,100_ closely follows the trend of the other repetitiveness measures, showing significant increases that are consistent with the compared measures. Further, we note that the relative increase in *π*_10,100_ is more prominent than in the other measures. This suggests a finer sensitivity of *π* when adding new information to the collection of data, making it a good candidate for detecting differences between genomes. We also confirmed the linear correlation between *π* and *r*, as observed by Oliva (Oliva, 2023). We performed this analysis using the same 60 *S. enterica* genomes from the previous experiment (see Figure 2).

**Figure 1.**
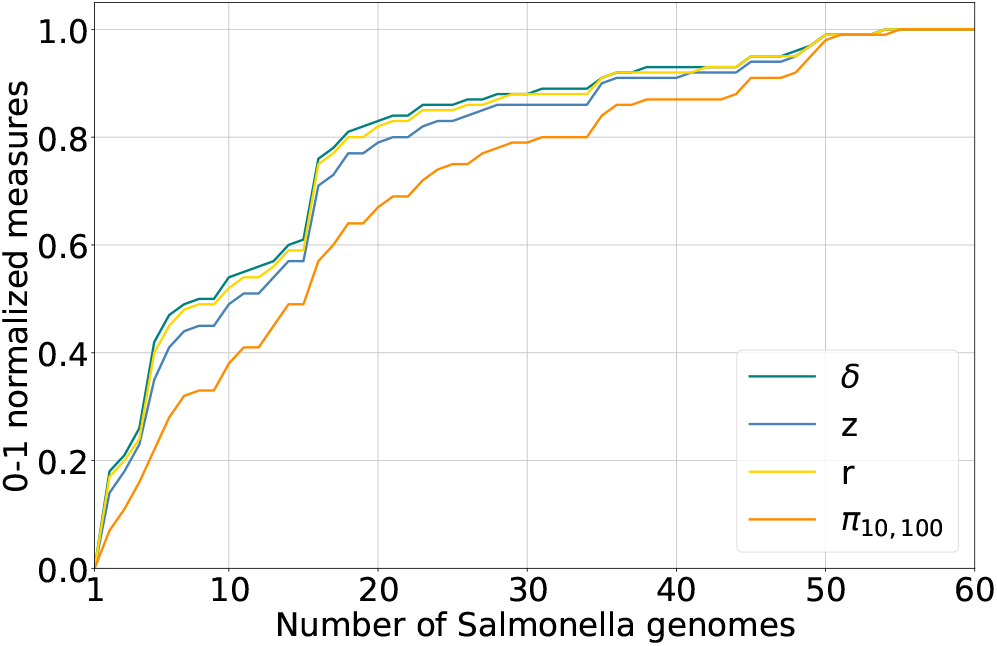
0-1 normalized measures. *δ, z, r* and *π*_10,100_ show a similar pattern of growth with the progressive addition of 60 genomes.

**Figure 2.**
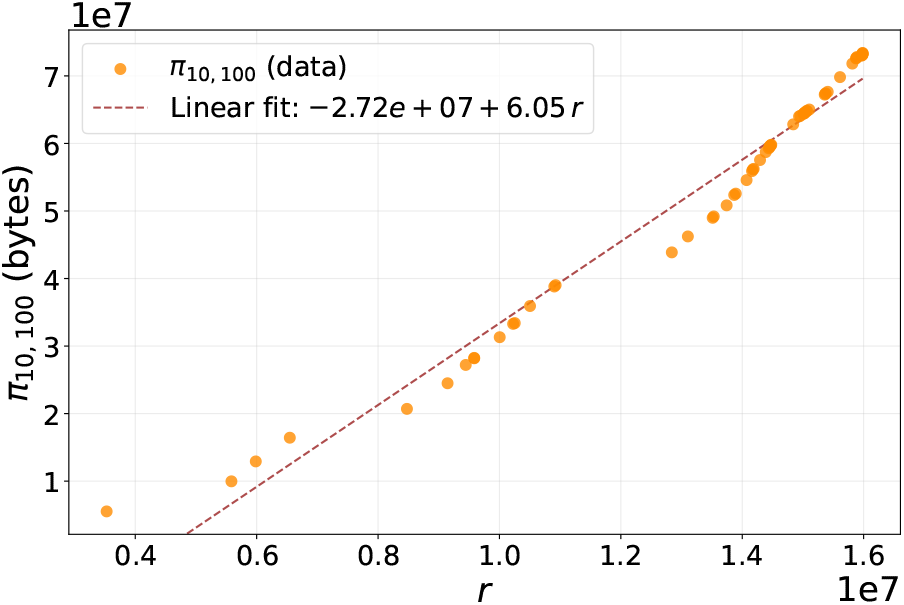
Linear correlation between *π*_10,100_ and r. The scatter plot illustrates individual data points, while the fitted line represents the linear relationship.

In a second set of experiments, we compared *π*_*w*,*p*_ of a random sequence to that of a real biological sequence of the same size to gain a better understanding of the value of *π*_*w*,*p*_ as a repetitiveness measure. We generated three different types of random sequences:

1. Each character is chosen according to the uniform distribution, i.e., with probability 1*/σ*.
2. Each character is chosen according to a background distribution, based on the character frequencies in the original sequence.
3. The sequence is a random permutation of the original sequence (C++ `shuffle`).

We report the results in Figure 3. As we can see, *π*_*w*,*p*_ of the random sequences is significantly larger than *π*_*w*,*p*_ of the real sequence.

**Figure 3.**
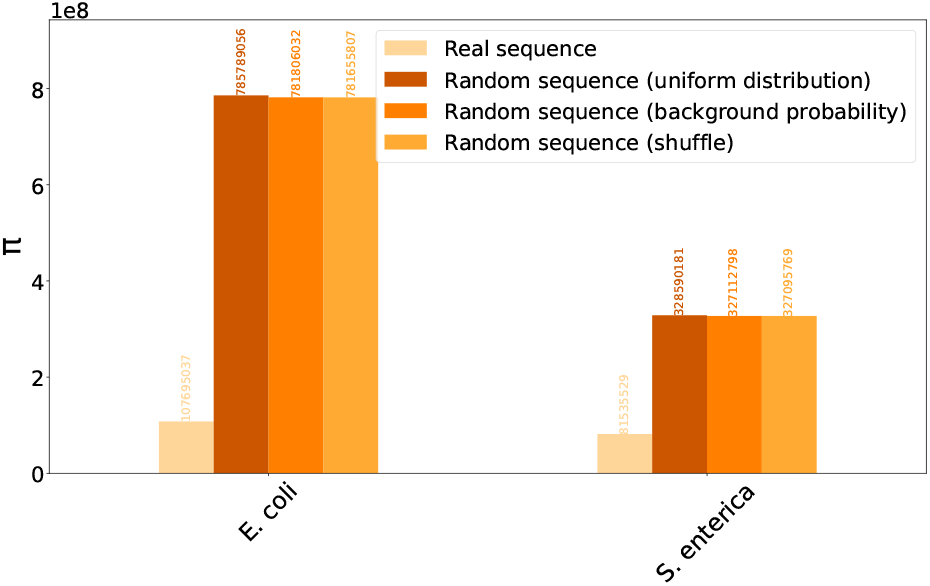
Comparison of *π*_*w*,*p*_ of a real sequence with a random sequence of the same size. The three random sequences are generated as described in Sec. 3.1

In a third set of experiments, we evaluated the efficiency of `piPFP` by comparing its running time and space consumption with those of other tools. Because we are now focusing on efficiency, we have added further tools to those used previously (namely, `PFP_LZ77`, `optimalBWT`, and `substring-complexity`). In particular we added: `kkp3`, `ropeBWT2`, `ropeBWT3`, `substring-complexity` (streaming version), and `DandD` (dashing mode). All tools used for the repetitiveness measure experiments are summarized in Table 2.

**Table 2:**
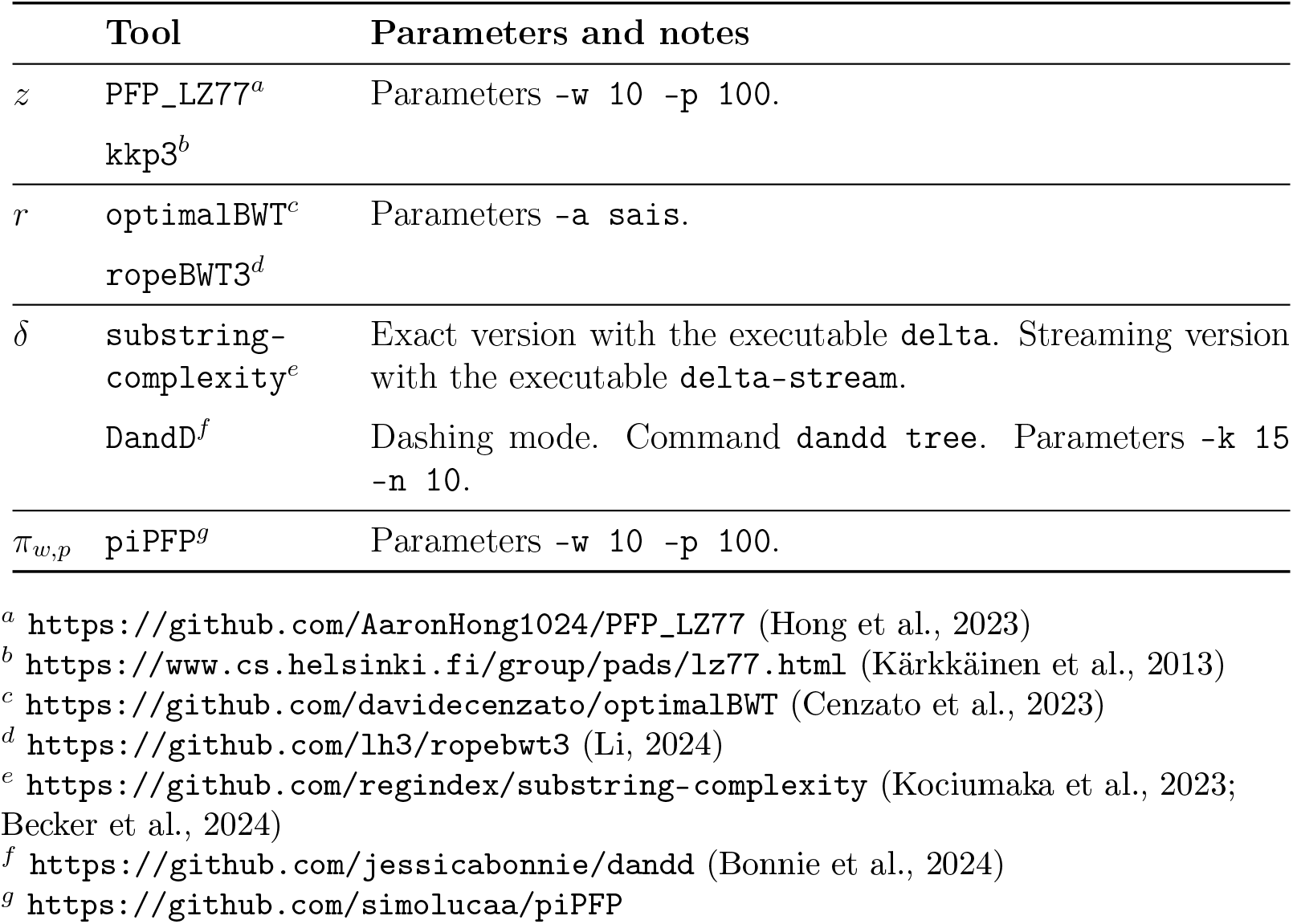
List of tools used for the repetitiveness experiments. The tools are grouped according to the measure they compute.

We ran every tool using a single thread on the full set of 60 *S. enterica* genomes and on another dataset consisting of 120 *E. coli* genomes. As shown in Figure 4a, our tool outperforms its competitors in terms of running time by at least an order of magnitude. It is worth noting that our tool also allows for high parallelization: using 16 threads, the time is reduced from 2.20 seconds to 0.34 seconds for *S. enterica* and from 5.34 seconds to 0.87 seconds for *E. coli*. Regarding space consumption, `piPFP` uses considerably less space than most of the other tools, except `DandD` and the streaming version of `substring-complexity`.

**Figure 4.**
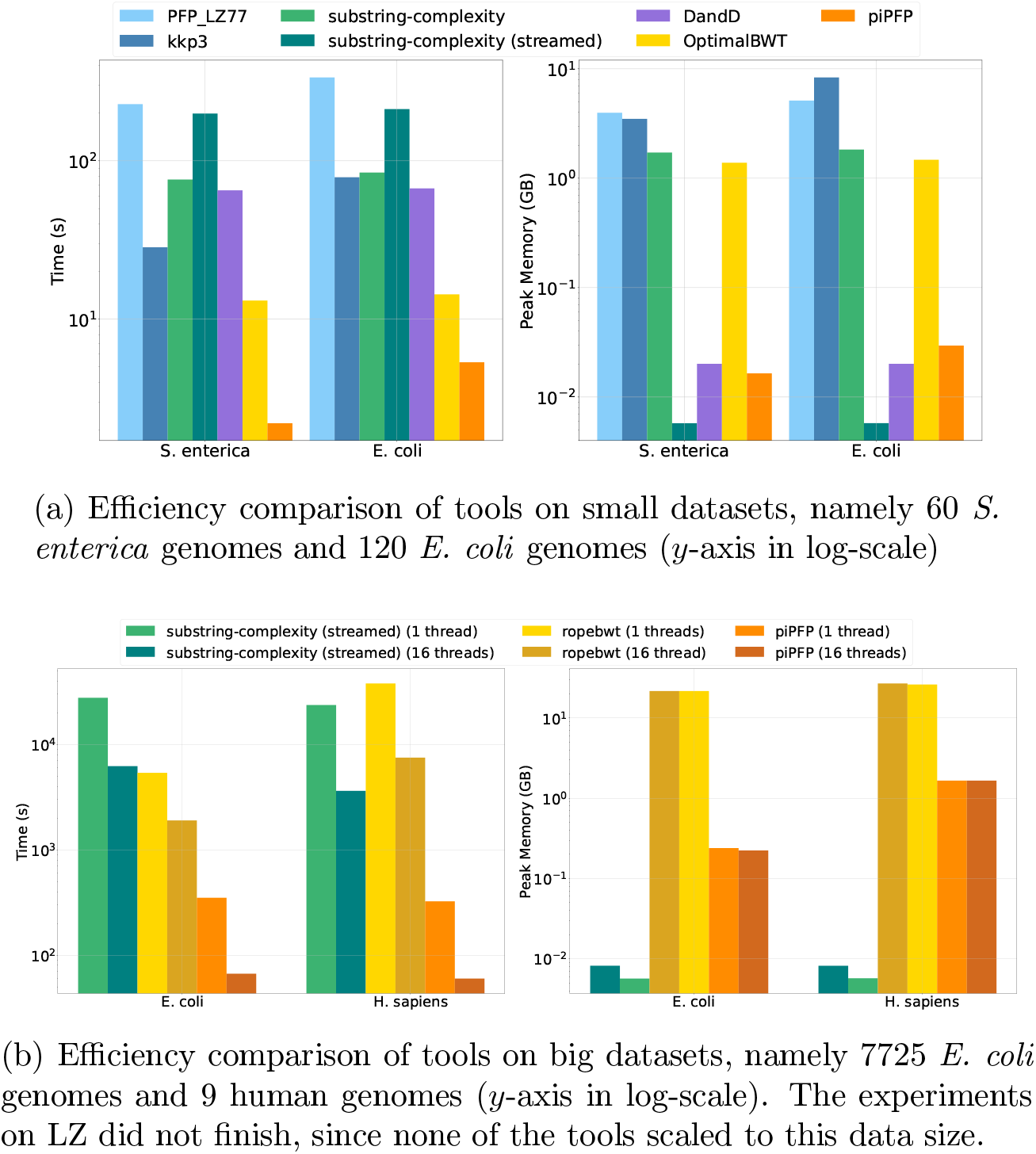
Efficiency comparisons on small and big datasets. The parameters choice for piPFP is *w* = 10 and *p* = 100.

### 3.2. PFP for estimating pangenome openness

To study the effectiveness of `piPFP` in estimating pangenome openness, we performed three sets of experiments. The first set involved a comparative analysis of `piPFP` against other tools. The second set focused on examining the impact of parameter selection on the accuracy of `piPFP`. The third set evaluated the computational efficiency of our tool. For the first two sets of experiments, we used the same datasets employed by Parmigiani et al. (Parmigiani et al., 2024). For the third set, we incorporated larger datasets, as detailed in Sec. 2.6.

In the first experiment, we analyzed 12 prokaryotic genomes (Table 1, entries 8-19) using three different tools: `Pangrowth`^2^(Parmigiani et al., 2024), `DandD` (Bonnie et al., 2024), and `piPFP` (our tool), with parameters (4, 10) and (5, 10) for (*w, p*).

In Table 3, we report the *α* values that indicate whether a pangenome is classified as open or closed. For completeness, we include the results from the gene-based tools reported in (Parmigiani et al., 2024) (Table 2 there): `Roary`^3^ (Page et al., 2015), `Pantools`^4^ (Jonkheer et al., 2022) and `BPGA`^5^ (Chaudhari et al., 2016). Out of the 12 datasets analyzed, `piPFP` classifies 10 as open, which aligns with the results from the other tools, while 2 are classified as closed. Specifically, the datasets designated as closed are *C. burnetii* (*α* = 1.44) and *Y. pestis* (*α* = 1.69). For *C. burnetii*, our tool’s classification agrees with `DandD` (*α* = 1.25), whereas for *Y. pestis*, our tool disagrees with all others.

**Table 3:** *α* values for piPFP, Pangrowth, DandD, Roary, Pantools and BPGA for each species.

| Species | #genes | piPFP<br>(4,10) | piPFP<br>(5,10) | Pangrowth | DandD | Roary | Pantools | BPGA |
| --- | --- | --- | --- | --- | --- | --- | --- | --- |
| *B. aphidicola* | 2911 | 0.22 | 0.24 | 0.30 | 0.25 | 0.20 | 0.21 | 0.20 |
| *B. cereus* | 295 | 0.68 | 0.71 | 0.78 | 0.79 | 0.68 | 0.68 | 0.69 |
| *C. botulinum* | 902 | 0.63 | 0.64 | 0.74 | 0.72 | 0.66 | 0.70 | 0.68 |
| *C. burnetii* | 2319 | 1.44 | 1.51 | 0.90 | 1.25 | 0.87 | 0.92 | 0.83 |
| *C. jejuni* | 242 | 0.84 | 0.87 | 0.87 | 0.90 | 0.76 | 0.78 | 0.69 |
| *F. tularensis* | 767 | 0.88 | 0.90 | 0.76 | 0.85 | 0.71 | 0.73 | 0.64 |
| *H. pylori* | 790 | 0.55 | 0.62 | 0.68 | 0.86 | 0.49 | 0.54 | 0.50 |
| *P. marinus* | 421 | 0.24 | 0.27 | 0.28 | 0.27 | 0.29 | 0.32 | 0.29 |
| *R. palustris* | 788 | 0.40 | 0.48 | 0.56 | 0.48 | 0.49 | 0.49 | 0.51 |
| *S. pneumoniae* | 818 | 0.85 | 0.88 | 0.75 | 0.78 | 0.73 | 0.77 | 0.69 |
| *S. pyogenes* | 674 | 0.76 | 0.78 | 0.75 | 0.76 | 0.81 | 0.74 | 0.68 |
| *Y. pestis* | 889 | 1.69 | 1.76 | 0.83 | 0.97 | 0.87 | 0.62 | 0.64 |

For this application, we chose small *w* and *p* for `piPFP` in order to get smaller phrases to capture the differences between genomes better. One might be tempted to view this as subsampling the *k*-mer space, for *k* = *p* + *w* −1. This is because, on average, we would expect every *p*th window to map to 0, resulting in roughly *n/p* phrases of length roughly *p* + *w* − 1. However, we found that the phrases from our experiments for small *w* and *p* result in significant variability of phrase lengths, which is far from a regular sampling of the *k*-mers present in the genomes (data not shown). It is intriguing to see that, nonetheless, the results we obtain with these parameters closely follow those from other tools.

In the second set of experiments, we studied whether the exact choice of parameters has a significant impact on the results. In Figure 5, we show that varying the parameters of `piPFP` does not have a noteworthy impact on the output *α* value. We tried every combination of *w* = {4, 5, 6, 7} and *p* = {10, 11, 12, 13, 14, 15}. As depicted in the boxplots, the distribution of the different *α* values has a very low variance. The standard deviation in all cases lay between 0.016 and 0.042 (see Table 4).

**Table 4:** Standard deviation and mean of *α* values.

| species | standard deviation | average $\alpha$ |
| --- | --- | --- |
| *S. pneumoniae* | 0.023 | 0.87 |
| *B. aphidicola* | 0.027 | 0.24 |
| *F. tularensis* | 0.026 | 0.89 |
| *P. marinus* | 0.039 | 0.28 |
| *S. pyogenes* | 0.017 | 0.78 |
| *C. jejuni* | 0.020 | 0.86 |
| *B. cereus* | 0.026 | 0.70 |
| *Y. pestis* | 0.042 | 1.75 |
| *H. pylori* | 0.040 | 0.61 |
| *C. burnetii* | 0.034 | 1.51 |
| *R. palustris* | 0.037 | 0.48 |
| *C. botulinum* | 0.025 | 0.64 |

**Figure 5.**
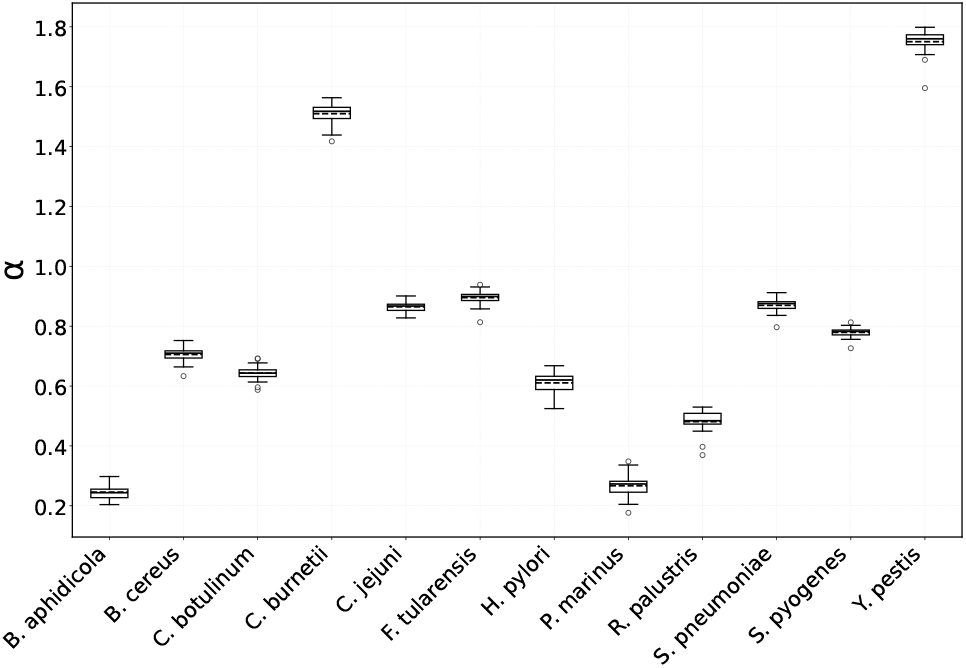
Distribution of *α* values on the 12 datasets. The values are taken over 24 parameter choices. See Sec. 3.2 for more details.

In the third set of experiments, we evaluated the efficiency of `piPFP` against `Pangrowth` and `DandD`. The results for `Pangrowth` and `piPFP` are shown in Figure 6. `DandD`‘s results are excluded because its computation time was significantly higher than that of the other two; this is due to its method for calculating pangenome growth, which involves averaging over a subset of permutations of genome unions. In this regard, we selected a subset of size 20, as the datasets are small and Bonnie et al. chose this parameter in the original DandD paper (Bonnie et al., 2024).

**Figure 6.**
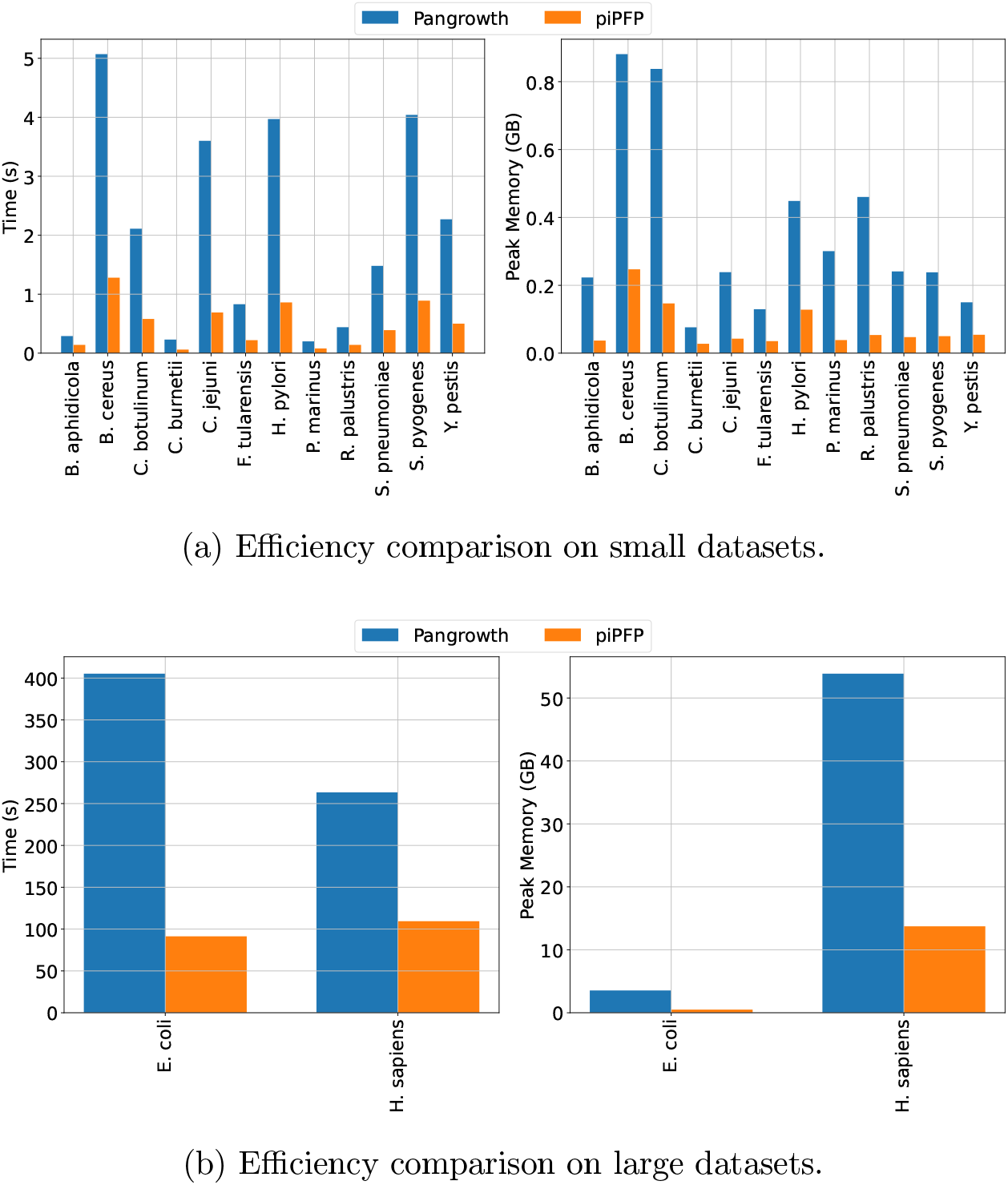
Efficiency comparison of Pangrowth and piPFP. The parameters choice for piPFP is *w* = 4 and *p* = 10. The maximum number of threads used by the two tools was set to 16.

**Figure 7.**
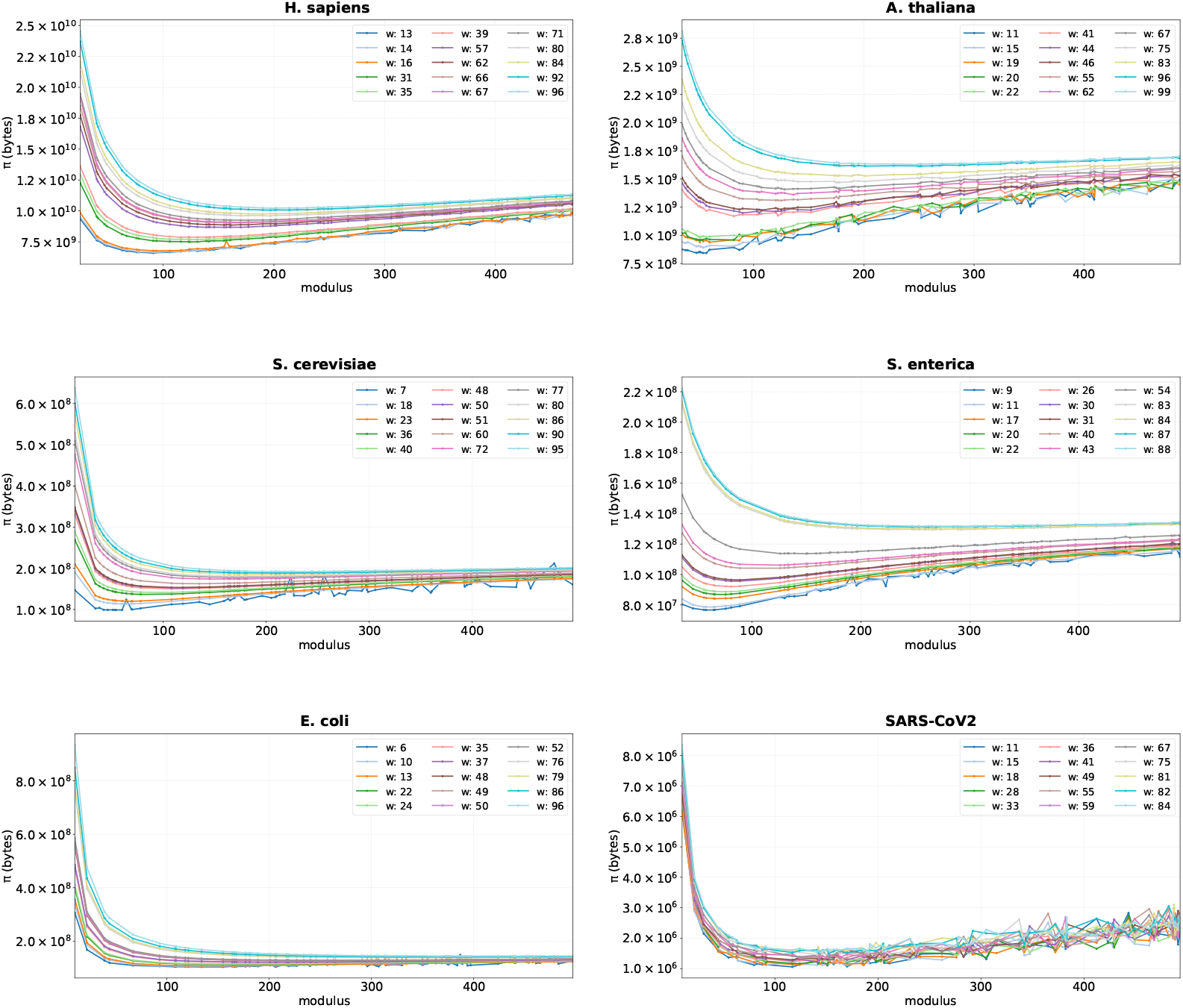
*π*_w,p_ with randomly chosen w and p. For each species, we chose 15 values for *w* from the range [4, 100] and 70 values for *p* from [10, 500].

In our experiments on smaller datasets (Table 1, entries 8-19), `DandD` consistently used less space compared to the other tools; however, the running times ranged from around 20 seconds to 45 minutes, depending on the dataset. While using a larger amount of space, our tool consistently stays below 1 GB of RAM, yet runs considerably faster. Compared to `Pangrowth`, `piPFP` consistently proved to be faster, used less space, and scaled well with the data (Fig. 6).

### 3.3. Parameters of PFP

The authors of the original paper, which introduced the PFP (Boucher et al., 2019), included a small study on the choice of the two parameters, the window size *w* and the modulus *p*. They gave experimental results on the Pizza and Chili repetitive corpus (Pizza and Chili repetitive corpus) with respect to compression (i.e., our *π*) with three parameter settings (6, 20), (8, 50), (10, 100), and on a biological dataset of 2.7 GB (1000 *S. enterica* genomes) with respect to running time and peak memory, with 15 combinations of parameters (*w* = 6, 8, 10 and *p* = 50, 100, 200, 400, 800). Based on their results, they chose the parameters (10, 100) as the default setting of their tool `BigBWT`. Oliva (Oliva, 2023) further studied how varying the parameters *w* and *p* affected ||*D*||, |*P* |, and the avarage length of a phrase. In his experiment, he tested combinations of *w* = 10, 15, …, 40 and *p* = 50, 100, 150, 200, 300, 400, 500, 600, 1200, 1800 on a dataset of 1000 diploid haplotypes of Chromosome 19, taken from the 1000 Genomes Project. Lastly, Lucà (Lucà, 2023) replicated the original study’s approach by testing the same 15 parameter combinations on three real datasets: *S. enterica, E. coli*, and SARS-CoV-2, alongside 36 artificial datasets derived from these real ones (which exhibited varying levels of repetitiveness).

We decided to study the choice of parameters more in depth. First, similarly to the other experiments, we chose regular intervals for our parameters, computing *π*_*w*,*p*_ for *w* = 5, 10, …, 30 and *p* = 40, 50, 60, …, 290, 300. The results showed an overall smooth function, in accordance with the results of previous studies. However, we observed a very irregular behavior with *w* = 5 (data not shown). Therefore, in a second set of experiments, we decided to use randomly chosen parameters to avoid artifacts due to possible number theoretic issues such as divisibility (see Sec. 3.3.1). On each of our six datasets, we computed *π*_*w*,*p*_ for over 1000 parameter combinations, choosing in each case 15 values for *w* from the interval [4, 100] and 70 values for *p* from the interval [10, 500]. We plot the size *π*_*w*,*p*_ of the resulting *PFP*(*T*) in Fig. 7. The values for *w* can be seen in the plots, while those for *p* can be found in the raw results file provided in the results GitHub repository (https://github.com/simolucaa/piPFP_experiments). We can see that in most cases, the graph for a fixed window size *w* and increasing modulus *p* shows an initial decrease, a minimum around *p* = 50 or *p* = 100, followed by a very gradual increase. This is because with modulus *p*, on average, there should be *n/p* phrases, where *n* is the total length of the dataset. Therefore, the parse will require 4*n/p* bytes, since in the parse, the phrases are referred to by their rank in the dictionary, and each integer requires 4 bytes. On the other hand, the dictionary will contain unique phrases only. If the sequences are very similar, a slight increase in phrase length should result in a slightly larger dictionary. These two effects result in a slight increase in the total size *π*_*w*,*p*_ for increasing *p*. Indeed, our experiments show that the parse length is very close to *n/p* for larger window sizes. (We discuss small window sizes separately below.) We observe that the SARS-CoV-2 data behaves more irregularly than the other datasets for larger window and modulus values. This is possibly due to the fact that viruses mutate frequently, and therefore, their genomes have more differences compared to those of other species.

To deepen this study, we also analyzed the behavior of *π* with fixed *p* and found that *π* increases with increasing *w* (data not shown). This may be due to the fact that increasing the parameter *w* increases the size of the overlaps between subsequent phrases. Indeed, if we assume that all phrases are unique, then |*P*| will be approximately *n/p*, independent of *w*, while the total size of the dictionary is ||*D*_*w*,*p*_|| = |*T* ^′^| + *w*(|*D*_*w*,*p*_| − 1), with the last term accounting for the fact that the overlaps are repeated between consecutive phrases. Therefore, if the number of phrases remains the same, then with increasing *w*, we get an increase in the total size of the overlaps, and consequently of ||*D*_*w*,*p*_||. Our experimental results indicate that a similar argument holds even if not all phrases are unique.

#### 3.3.1. Small window sizes

As can be seen in several of the plots in Fig. 7, there appears to be an irregular behavior of the size of the data structure for small window sizes. This is because for certain values of *p*, there are no or very few trigger strings, resulting in one long phrase, and thus no compression at all. This phenomenon can be seen much more in detail in Fig. 8, where we plot, for the same six datasets, the measure *π*_*w*,*p*_ using window sizes *w* = 4, 5, …, 10 and modulus *p* taking values of multiples of 10 between 40 and 300. For example, on *S. enterica*, there are no trigger strings in the text at all for *w* = 5 and *p* = 160, 170, or for *w* = 4 and *p* = 40, 80, 120, 160, 170, 200, 230, 240, 250, 280, resulting in one long phrase.

**Figure 8.**
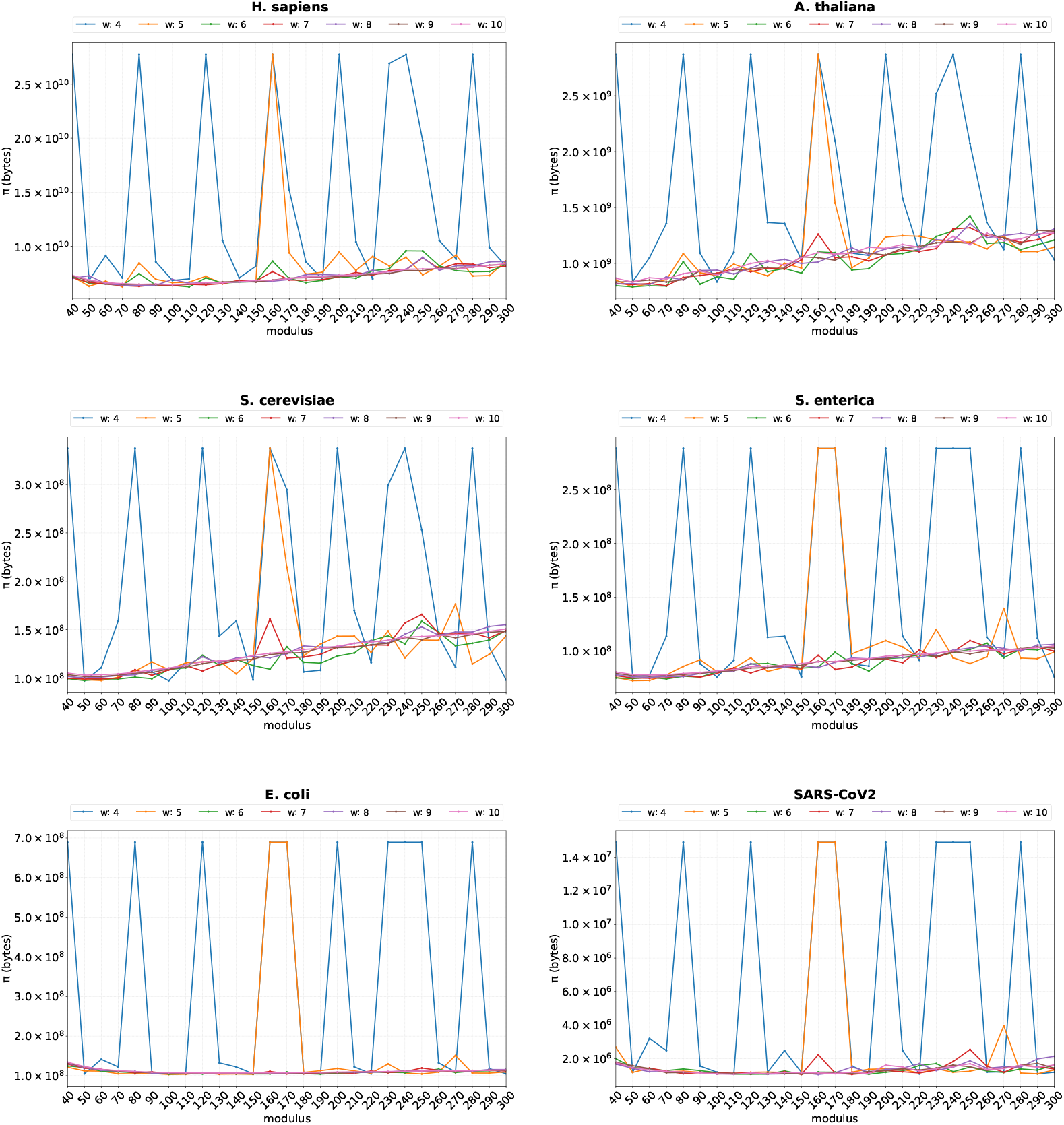
Values of *π*_w,p_ with small w. This highlights the unstable behavior of *π* with small *w*, especially with *w* = 4, 5.

This effect is due to a phenomenon which we refer to as *coefficient scarcity*. Since the alphabet is very small, and the characters are encoded according to their ASCII-code, it can happen that no string of size *w* is hashed to 0 modulo *p*. For example, if *p* is a multiple of 40, then no string of length 4 over the alphabet {`A`, `C`, `G`, `T`} maps to 0 modulo *p*, as we show next.

Let Σ = {`A`, `C`, `G`, `T`}, *w* = 4, and let *p* be a multiple of 40. Consider a string *S* = *s*_1_*s*_2_*s*_3_*s*_4_ ∈ Σ^4^, and define *x*_*i*_ = *f* (*s*_*i*_), the ASCII-256 encoding of character *s*_*i*_. In particular,

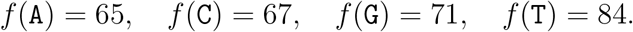

Let *b* = 256 be the base. Then *S* is a trigger string if and only if

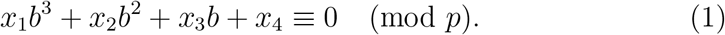

Since *p* is a multiple of 40, Eq. (1) implies

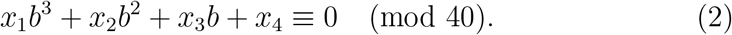

As can be easily verified, *b, b*^2^, *b*^3^ are all congruent 16 (mod 40), and therefore, Eq. (2) equation reduces to

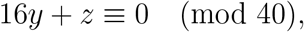

where *y* = *x*_1_ + *x*_2_ + *x*_3_ (mod 40) and *z* = *x*_4_ (mod 40). Now note that the possible values for *z* are 25, 27, 31, and 4, as these are the values of *f* (*c*) (mod 40) for the four characters *c* = `A`, `C`, `G`, `T`. Since 16*y* is the additive inverse modulo 40 of *z*, it must therefore be equal to one of 15, 13, 9, or 36. However, the set generated by the element 16 in ℤ_40_ is ⟨16⟩ = {0, 16, 32, 8, 24}, and, in particular, does not contain any of those numbers. This proves that Eq. (1) has no solution.

Note also that we could ignore the large prime *q* used for the Karp-Rabin hash in this argument because with such small window sizes, all numbers in question are smaller than *q*.

As can be seen from Fig. 7, the coefficient scarcity effect disappears with increasing window size.

To alleviate this phenomenon, one could consider changing the corresponding values to which the individual characters are mapped, such as mapping them to random or prime numbers. However, in most implementations, the standard ASCII encoding is used by default.

## 4. Conclusion

In this paper, we presented *π*, the size of the PFP data structure, and proposed it as a measure of repetitiveness of the text on the one hand, and as a method for deciding pangenome openness, on the other. For both uses, we showed that our new measure gives comparable results to other, more established, measures. We also presented a tool, `piPFP`, computing it, and showed that it outperforms other tools both w.r.t. running time and space. To be fair, our tool is tailor-made to compute only *π*, while several (but not all) of the other methods compute the measures only as a by-product. Nonetheless, we would argue that it makes sense to use a more efficient tool if one only needs the value itself.

On the theoretical side, our experiments show that the PFP heuristic does a good job in capturing the string’s repetitiveness, similar to more structured (and well-defined) measures such as *z, r*, or *δ*. From our study of the pangenome openness, we conclude that PFP can also be used successfully to simulate *k*-mer counts.

It remains an open problem to compute *π*^*^(*T*), the minimum size of the PFP of the text *T*, taken over all choices of parameter pairs (*w, p*). Our results from Section 3.3 can serve as a starting point here. In particular, we saw that while the coefficient scarcity problem seems to be restricted to small window sizes for most of our datasets, on the SARS-CoV-2 data, we observed an irregular behavior for all window sizes, possibly due to the fact that this data set, being a viral genome, is far more variable than the others. This seems to imply that on more variable data, the choice of parameters has a higher impact on *π*.

Future work includes exploring further applications of *π*, such as for pairwise string similarity resp. distance, i.e., as a non-alignment based string similarity measure. Finally, we believe that studying the dictionary size alone could be of interest, especially as a measure of inter-sequence similarity within a dataset of pairwise similar sequences, such as a pangenome.

## 5. Acknowledgments

ZsL is partially funded by the MUR PRIN project no. 2022YRB97K ‘PINC’ (Pangenome INformatiCs: From Theory to Applications), and by the INdAM-GNCS project no. E53C24001950001.

## Footnotes

1 https://gitlab.com/manzai/Big-BWT

2 https://github.com/gi-bielefeld/pangrowth

3 https://github.com/sanger-pathogens/Roary

4 https://git.wur.nl/bioinformatics/pantools

5 https://sourceforge.net/projects/bpgatool/

